# Prediction of Lignocellulose Degradation Potential of Wood Decay Fungi using Comparative Genomic Analysis

**DOI:** 10.64898/2026.06.15.732456

**Authors:** Siri Tantry, Steven Ahrendt, Guifen He, Kurt LaButti, Anna Lipzen, Kerrie Barry, David Culley, Jon Magnuson, Joseph Spatafora, Igor V. Grigoriev

## Abstract

The Agaricomycotina accounts for roughly a third of all described fungi. They are important due to their wide range of lifestyles and economic and environmental relevance. Certain agaricomycetes act as lignocellulose degraders, playing a significant role in forest ecosystems and bioremediation processes.

These wood-decaying fungi have historically been classified as mostly white- or brown-rot based on their ability to degrade lignin, with white-rot fungi possessing a collection of lignocellulose-degrading enzymes, which are reduced or absent in brown-rot fungi. Here, we sequenced and annotated the genome of the agaricomycete *Crepidotus cesatii* CBS 511.95 and explored its genome and predicted enzymatic content in a comparative context. The 36.04 Mbp genome is in 235 scaffolds, with 3.34% repeat content and 12,891 predicted genes. We found that the PFAM distributions of identified orthogroups suggested that *C. cesatii* shows patterns more similar to white-rot fungi compared to brown-rot fungi. Additionally, *C. cesatii* contained multiple copies of CAZymes CBM1 and AA9 involved in hydrolysis of lignocellulose, similar to white-rot fungi. On the other hand, according to the Conserved Unique Peptide Patterns (CUPP) data for AA2 peroxidases, the key enzymes in lignin degradation, *C. cesatii* is more similar to brown-rot fungi. Based on our analyses we predict that *C. cesatii* is another representation of the continuum of wood decaying modes between white and brown rot fungi combining genetic features of both types of fungi.

## Introduction

The fungal subphylum Agaricomycotina, one of three within the Basidiomycota, accounts for 34% of described fungi. While perhaps best known for containing mushroom-forming species, the Agaricomycotina contain a diversity of fungal morphologies and are important to study in a variety of contexts, including culinary (containing edible mushrooms), economic (containing pathogens of various crops), and medical (containing human pathogens) (De Mattos-Shipley et al., 2016). Additionally, certain Agaricomycotina fungi can act as natural lignocellulose degraders, playing important roles in forest ecosystems, biogeochemical cycles, biodegradation, and bioremediation processes.

Such wood decay fungi have traditionally been classified as either brown or white rot, based on their abilities to decompose constituent plant cell wall compounds: cellulose, hemicellulose, and lignin. Brown rot fungi use non-enzymatic activity (such as Fenton chemistry) to degrade cellulose and hemicellulose while leaving lignin modified but largely intact (Veloz Villavicencio et al., 2020). White rot fungi by contrast, enzymatically degrade cellulose, hemicellulose, and lignin (Floudas et al., 2012; Martínez et al., 2005). For lignin degradation, white-rot fungi tend to have a high number of class II peroxidase (AA2) genes, encoding lignin peroxidases (LiPs), manganese peroxidases (MnPs), or versatile peroxidases (VPs), absent in brown-rot fungi (Veloz Villavicencio et al., 2020). Additionally, enzymatic H_2_O_2_ generation has been observed primarily in white rot fungi using flavoproteins (AA3) and copper radical oxidases (AA5), while is very limited or nonexistent in brown-rot fungi (Hasegawa et al., 2024). Other important enzymes employed by white-rot fungi include cellulose-binding (CBM1) domain proteins and lytic polysaccharide monooxygenases (AA9) (Ruiz-Dueñas et al., 2021).

While traditional binary classification of wood decay fungi relied on a selection of functionally characterized enzymes, the complete picture of fungal lignocellulose degradation has not been fully understood. Previous work identified fungi capable of degrading lignin despite lacking the traditionally associated class II peroxidase enzymes (but sharing cellulose binding proteins and monooxygenases), suggesting a continuum of nutritional modes and intermediate (“uncertain”) wood decay strategies. Such modes are exhibited by fungi like *Botryobasidium botryosum*, *Schizophyllum commune*, and *Jaapia argillacea* (Riley et al., 2014). Furthermore, the complete set of enzymes may involve numerous gene families with as-yet undetermined functions (Nagy et al., 2017). Deep sequencing and genome analyses of species within the Agaricomycotina will augment our understanding of the complexities of their lignocellulose-degradation strategies. Here, we describe the sequencing, annotation, and characterization of the genome of *Crepidotus cesatii* CBS 511.95 (family Crepidotaceae, order Agaricales). Only one genome from the Crepidotaceae family is currently available, that of *Crepidotus variabilis* CBS 506.95 (Ruiz-Dueñas et al., 2021). While it is reported to be one of the most efficient basidiomycetes in lignin degradation with one of the highest cellulose enrichments (Martínez et al. 2005), its growth on forest wood debris is considered uncharacterized (Ruiz-Dueñas et al., 2021). Field surveys in the UK suggest that *C. cesatii* and *C. variabilis* can both be considered commonly occurring fungi (Fortey, 2017). The genome analysis of *C. cesatii* will help elucidate not only the specific decay mechanisms of the Crepidotaceae family, but also the more general patterns observed across the Agaricomycotina.

## Methods

### Growth and DNA extraction

*Crepidotus cesatii* CBS 511.95 was grown in shake flask liquid cultures at room temperature and 30⁰C. Filtered biomass was ground with a mortar and pestle in liquid nitrogen. Genomic DNA and RNA were both extracted with Trizol. The DNA was treated with RNase, and the RNA was reprecipitated with LiCl.

### Genome sequencing and assembly

The *Crepidotus cesatii* CBS 511.95 genome sequencing library was prepared as follows: 4732 ng of genomic DNA was sheared to >10kb using Covaris g-Tubes. The sheared DNA was treated with exonuclease to remove single-stranded ends and DNA damage repair mix followed by end repair and ligation of blunt adapters using SMRTbell Template Prep Kit 1.0 (Pacific Biosciences). The library was purified with AMPure PB beads. PacBio Sequencing primer was then annealed to the SMRTbell template library and Version P6 sequencing polymerase was bound to them. The prepared SMRTbell template libraries were then sequenced on a Pacific Biosciences RSII sequencer using Version C4 chemistry and 1x240 sequencing movie run times.

After sequencing, filtered subread data was assembled together with Falcon version 0.7.3 (https://github.com/PacificBiosciences/FALCON) to generate an initial assembly. Mitochondria were assembled separately from the Falcon pre-assembled reads (preads) using an in-house tool (assemblemito.sh), used to filter the preads, and polished with Quiver version smrtanalysis_2.3.0.140936.p5 (https://github.com/PacificBiosciences/GenomicConsensus). A secondary Falcon assembly was generated using the mitochondria-filtered preads, improved with finisherSC version 2.0 (Lam et al., 2015), and polished with Quiver.

### Transcriptome sequencing and assembly

A plate-based RNA sample prep was performed on the PerkinElmer Sciclone NGS robotic liquid handling system using Illumina’s TruSeq Stranded mRNA HT sample prep kit utilizing poly-A selection of mRNA following the protocol outlined by Illumina in their user guide: https://support.illumina.com/sequencing/sequencing_kits/truseq-stranded-mrna.html, and with the following conditions: total RNA starting material was 1000 ng per sample and 8 cycles of PCR was used for library amplification. Sequencing was performed using an Illumina instrument. Using BBDuk (https://sourceforge.net/projects/bbmap/), raw reads were evaluated for artifact sequence by kmer matching (kmer=25), allowing 1 mismatch, and the detected artifact was trimmed from the 3’ end of the reads. RNA spike-in reads, PhiX reads, and reads containing any Ns were removed. Quality trimming was performed using the phred trimming method set at Q6. Finally, following trimming, reads under the length threshold were removed (minimum length 25 bases or 1/3 of the original read length - whichever is longer). Filtered reads were assembled into consensus sequences using Trinity ver. 2.3.2 (Grabherr et al., 2011) with the --normalize_reads and --jaccard_clip options. For additional information, see: http://trinityrnaseq.github.io.

### Genome annotation

The nuclear genome was annotated using the JGI Fungal Annotation Pipeline, which utilizes the assembled transcriptome and combines several gene prediction and annotation methods (Grigoriev et al., 2014; Kuo et al., 2014). The mitochondrial genome was annotated separately (Haridas et al., 2018). For targeted functional analyses, we used predicted PFAM domains (Mistry et al., 2021), CAZymes (Carbohydrate Active Enzymes - Drula et al., 2022), and CUPP analysis (Conserved Unique Peptide Patterns - Barrett et al., 2020).

### Phylogenomic and Functional Analysis

For comparative genomics, our dataset (Table 1) consisted of 38 published fungal genomes available in MycoCosm (Grigoriev et al., 2014). ORTHOFINDER v3.0 (Emms & Kelly, 2019) was used to identify single-copy orthologs, which were aligned using MAFFT v7 (Katoh & Standley, 2013). The alignments were trimmed using CLIPKIT v2.3.0 (Steenwyk et al., 2020) and used to build a phylogenetic tree using FastTree v2 (Price et al., 2010). We visualized the tree using the R Phytools package (Revell, 2024).

**Table 1:** Genome comparison of *C. cesatii* and its phylogenetically close *C. variabilis* along with other fungi. *BUSCO scores represent single-copy counts using the fungi_odb10 lineage dataset

| Species Name | MycoCosm ID | Size (Mbp) | Repeats (Mbp) | # Genes | G+C, % | BUSCO, %* | Publication |
| --- | --- | --- | --- | --- | --- | --- | --- |
| *Agaricus bisporus* var. *bisporus* (H97) | Agabi_varbisH97_2 | 30.23 | 0.66 | 10438 | 46.5 | 93.8 | Morin et al., 2012 |
| *Aspergillus nidulans* FGSC A4 | Aspnid1 | 30.48 | 0.95 | 10680 | 50.4 | 98.3 | Galagan et al., 2005 |
| *Auricularia subglabra* | Aurde3_1 | 76.85 | 1.46 | 25459 | 58.6 | 96.3 | Floudas et al., 2012 |
| *Botryobasidium botryosum* | Botbo1 | 46.67 | 3.13 | 16526 | 52.4 | 98.5 | Riley et al., 2014 |
| *Ceriporiopsis subvermispora* B | Cersu1 | 38.97 | 0.39 | 12125 | 53.8 | 90 | Fernandez-Fueyo et al., 2012 |
| *Coniophora puteana* | Conpu1 | 42.97 | 0.61 | 13761 | 52.4 | 93.4 | Floudas et al., 2012 |
| *Coprinopsis cinerea* | Copci1 | 36.29 | 0.77 | 13393 | 51.7 | 95.6 | Stajich et al., 2010 |
| *Crepidotus cesatii* CBS 511.95 | Creces1 | 36.04 | 1.2 | 12891 | 48 | 96.8 | This study |
| *Crepidotus variabilis* CBS 506.95 | Crevar1 | 38.58 | 0.93 | 14573 | 46.8 | 98.9 | Ruiz-Dueñas et al., 2021 |
| *Cryptococcus neoformans* var. *grubii* H99 | Cryne_H99_1 | 18.87 | 0.16 | 6967 | 48.2 | 95 | Janbon et al., 2014 |
| *Dacryopinax primogenitus* DJM 731 SSP1 | Dacsp1 | 29.5 | 0.78 | 10242 | 52.2 | 93.4 | Floudas et al., 2012 |
| *Dichomitus squalens* LYAD-421 SS1 | Dicsq1 | 42.75 | 1.76 | 12290 | 55.6 | 95.4 | Floudas et al., 2012 |
| *Fomitiporia mediterranea* | Fomme1 | 63.35 | 23.53 | 11333 | 40.8 | 95.3 | Floudas et al., 2012 |
| *Fomitopsis schrenkii* FP-58527 SS1 | Fompi3 | 41.61 | 0.54 | 13885 | 55.7 | 98 | Floudas et al., 2012 |
| *Galerina marginata* | Galma1 | 59.42 | 1.18 | 21461 | 48 | 97.6 | Riley et al., 2014 |
| *Gloeophyllum trabeum* | Glotr1_1 | 37.18 | 1.16 | 11846 | 53.2 | 98.7 | Floudas et al., 2012 |
| *Heterobasidion annosum* | Hetan2 | 33.65 | 1.28 | 13383 | 52.2 | 97.1 | Olson et al., 2012 |
| *Jaapia argillacea* | Jaaar1 | 45.05 | 2.11 | 16419 | 49.8 | 98 | Riley et al., 2014 |
| *Laccaria bicolor* | Lacbi2 | 60.71 | 15.06 | 23132 | 46.9 | 97.2 | Martin et al., 2008 |
| *Malassezia globosa* | Malgl1 | 8.96 | 0.07 | 4286 | 52.1 | 76.8 | Xu et al., 2007 |
| *Melampsora larici-populina* | Mellp2_3 | 109.88 | 49.65 | 19550 | 41 | 87.6 | Persoons et al., 2022 |
| *Phanerochaete carnos*<br>HHB-10118-Sp | Phaca1 | 46.29 | 2.81 | 13937 | 53.1 | 87.7 | Suzuki et al., 2012 |
| *Phanerochaete chrysosporium* RP-78 | Phchr2 | 35.15 | 1.96 | 13602 | 56.8 | 98 | Ohm et al., 2014 |
| *Piriformospora indica*<br>DSM 11827 from MPI | Pirin1 | 24.98 | 0.73 | 11767 | 50.7 | 92.6 | Zuccaro et al., 2011 |
| *Pleurotus ostreatus* PC15 | PleosPC15_2 | 34.34 | 0.58 | 12330 | 50.9 | 94.7 | Alfaro et al., 2016 |
| *Postia placenta*<br>MAD-698-R-SB12 | PosplRSB12_1 | 42.45 | 5.54 | 12541 | 53.8 | 95.4 | Gaskell et al., 2017 |
| *Puccinia graminis* f. sp.<br>*tritici* | Pucgr2 | 88.64 | 16.91 | 15979 | 43.4 | 79.7 | Cuomo et al., 2017 |
| *Punctularia strigosozonata* | Punst1 | 34.17 | 0.8 | 11538 | 55.2 | 94.6 | Floudas et al., 2012 |
| *Schizophyllum commune*<br>H4-8 | Schco3 | 38.67 | 1.35 | 16319 | 57.5 | 96.8 | Marian et al., 2022 |
| *Serpula lacrymans* S7.9 | SerlaS7_9_2 | 42.73 | 12.97 | 12789 | 45.3 | 98.4 | Eastwood et al., 2011 |
| *Stagonospora nodorum*<br>SN15 | Stano2 | 37.21 | 1.07 | 12380 | 50.4 | 93 | Hane et al., 2007 |
| *Stereum hirsutum*<br>FP-91666 SS1 | Stehi1 | 46.51 | 0.59 | 14072 | 51.3 | 95.5 | Floudas et al., 2012 |
| *Trametes versicolor* | Trave1 | 44.79 | 0.48 | 14296 | 57.7 | 95.8 | Floudas et al., 2012 |
| *Tremella mesenterica*<br>Fries | Treme1 | 28.64 | 0.66 | 8313 | 46.7 | 94.7 | Floudas et al., 2012 |
| *Trichoderma reesei* QM6a | Trire2 | 33.45 | 0.85 | 9143 | 52.8 | 99.3 | Martinez et al., 2008 |
| *Ustilago maydis* 521 | Ustma2_2 | 19.66 | 0.37 | 6785 | 54 | 97.8 | Kämper et al., 2006 |
| *Wallemia mellicola* CBS<br>633.66 | Walse1 | 9.82 | 0.07 | 5284 | 40 | 92.3 | Padamsee et al., 2012 |
| *Wolfiporia cocos* MD-104<br>SS10 | Wolco1 | 50.48 | 5.06 | 12746 | 52.2 | 98.2 | Floudas et al., 2012 |

## Results and Discussion

### Genome Analysis and Phylogeny

The *Crepidotus cesatii* genome was assembled into 235 contigs, with a total assembly size of 36.04 Mbp (N50=23; L50=0.51 Mbp), GC content of 47.97%, and 3.34% repeats. The largest contig was 1.25 Mbp. The assembly size falls within the range of phylogenetically related organisms such as *Crepidotus variabilis* and other Agaricomycotina such as *Serpula lacrymans* and *Schizophyllum commune* (Fig.1). Additionally, *C. cesatii* is slightly smaller than the 38.58 Mbp genome of *Crepidotus variabilis* CBS 506.95.

**Figure 1:**
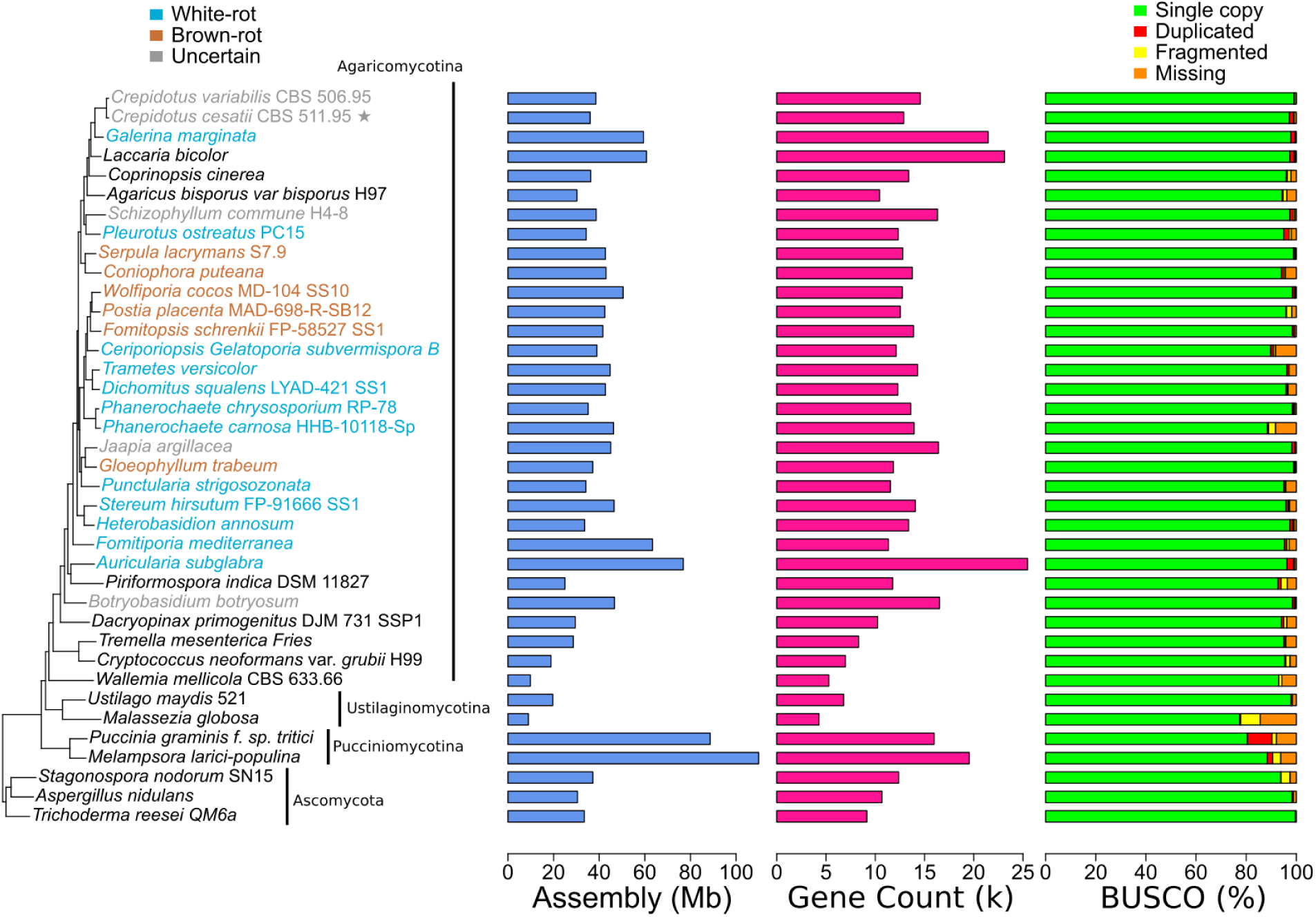
Phylogenetic tree with assembly size, gene counts, and completeness (BUSCO) for a set of Basidiomycetes with Ascomycete outgroups. The genome of interest, *Crepidotus cesatii* is indicated with a star (*).

The nuclear genome contained 12,891 predicted protein-coding genes, with 57.26% encoding known protein PFAM domains. Genome completeness was estimated at 99.78% using CEGMA (Parra et al., 2007) and 97.2% using BUSCO (Simão et al., 2015) fungi_odb10 lineage dataset. Similar to *Crepidotus variabilis*, *C. cesatii* displayed high BUSCO completeness (completeness classified when lengths are within two standard deviations of the BUSCO group mean length) and similar gene count and assembly length (Table 1).

The mitochondrial genome was assembled into a single circular scaffold of 78.1 kb and 27% GC content. All 15 core fungal mitochondrial genes were identified, including 14 oxidative phosphorylation subunit genes (*atp6, atp8, atp9, cob, cox1, cox2, cox3, nad1,nad2, nad3, nad4, nad4L, nad5, nad6*), as well as *rps3*. Both large and small rRNA subunits were predicted, along with 25 tRNAs covering all amino acids. By comparison, while the *C. variabilis* mitochondrial genome similarly contains 15 core protein coding genes, 25 tRNA genes, and both rRNA subunits, it is in a smaller scaffold of 42.2 kb. The *C. cesatii* genes collectively contained 19 introns, compared to only 5 found in *C. variabilis*. Correspondingly, *C. cesatii* shows more homing endonuclease genes (15 LAGLIDADG, 1 GIY_YIG) compared to *C. variabilis* (1 LAGLIDADG, 1 GIY_YIG).

Using OrthoFinder, we identified 404 single-copy ortholog clusters in our dataset of 38 wood-decaying and other fungi. We constructed a phylogenetic tree based on the alignment of these orthologs where the bargraphs visually represented assembly in Mbp, Gene Counts, as well as BUSCO completeness (including representation of single-copy, duplicated, fragmented, and missing BUSCO categories).

Similar to the tabular genome characteristics, the bargraphs also visually illustrate that *C. cesatii* and *C. variabilis* are similar in terms of assembly, gene counts, and BUSCO completeness (Figure 1).

### CAZyme and CUPP Predictions

To better understand and compare the fungal metabolic processes, we compared CAZyme annotations across different wood-decaying fungi to find unique patterns associated with the decay mode classification. In agreement with previous research (Hasegawa et al., 2024), we observed that white-rot fungi tended to have large numbers of genes in families AA2 (class II lignin-modifying peroxidases), AA3 (the glucose-methanol-choline (GMC) oxidoreductases), AA9 (monooxygenases of the Auxiliary Activity Family 9), and CBM1 (carbohydrate-binding module). In contrast, brown-rot fungi showed reduced counts of GH6 (Glycoside Hydrolase Family 6) and GH7 (Glycoside Hydrolase Family 7) families. The *C. cesatii* genome encoded multiple copies of CBM1, AA9, GH6, and GH7 proteins (Table 2), consistent with white-rot fungi, but only a single AA2 protein (protein ID 272474) typically abundant in white-rot fungi. This single AA2 protein distinguishes *C. cesatii* from white-rot fungi and deviates toward fungi with uncertain wood-decay types that share some features of both groups (Floudas et al., 2015; Riley et al., 2014). Interestingly, *C. variabilis* shows a similar pattern.

**Table 2:**
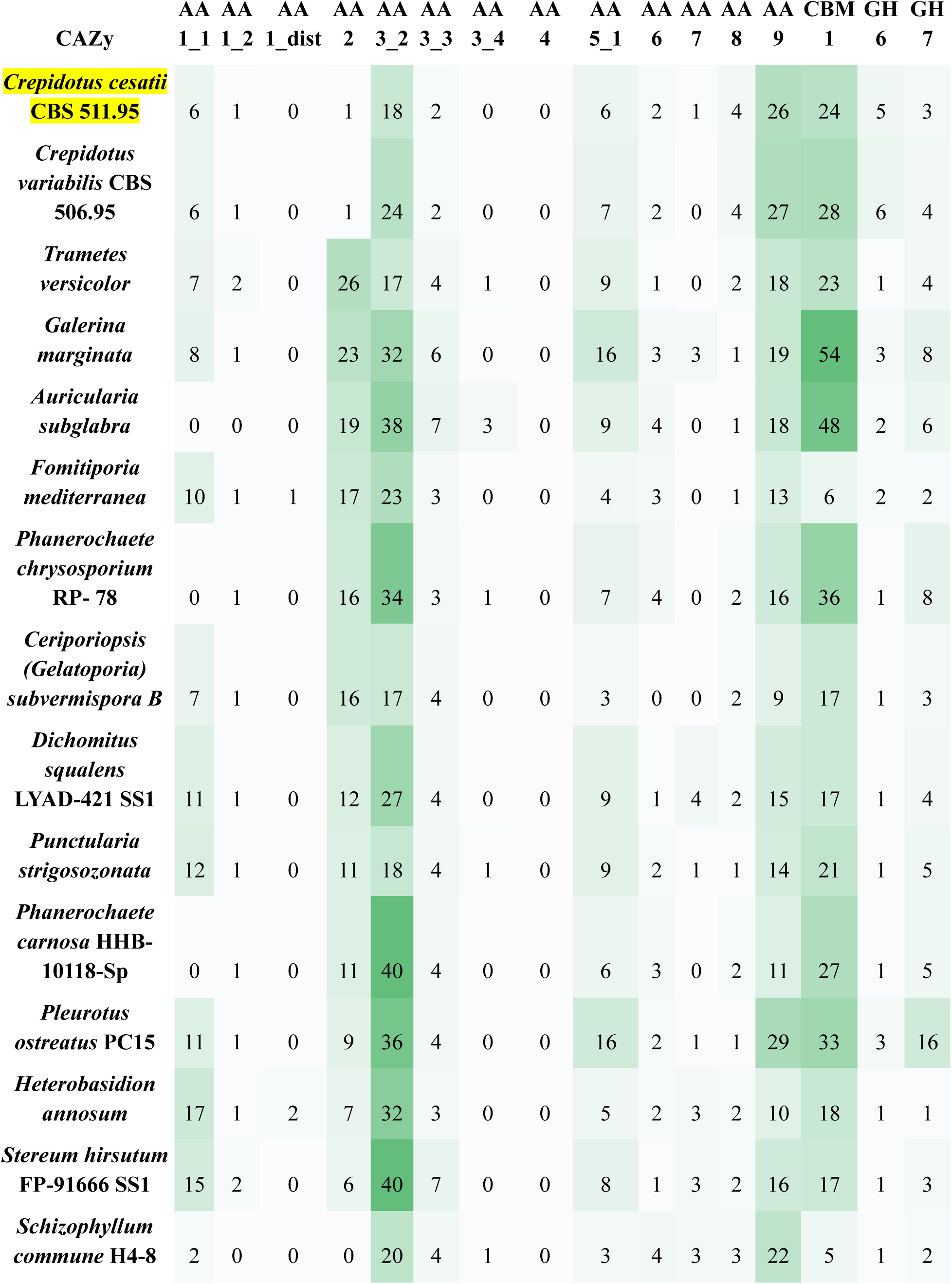
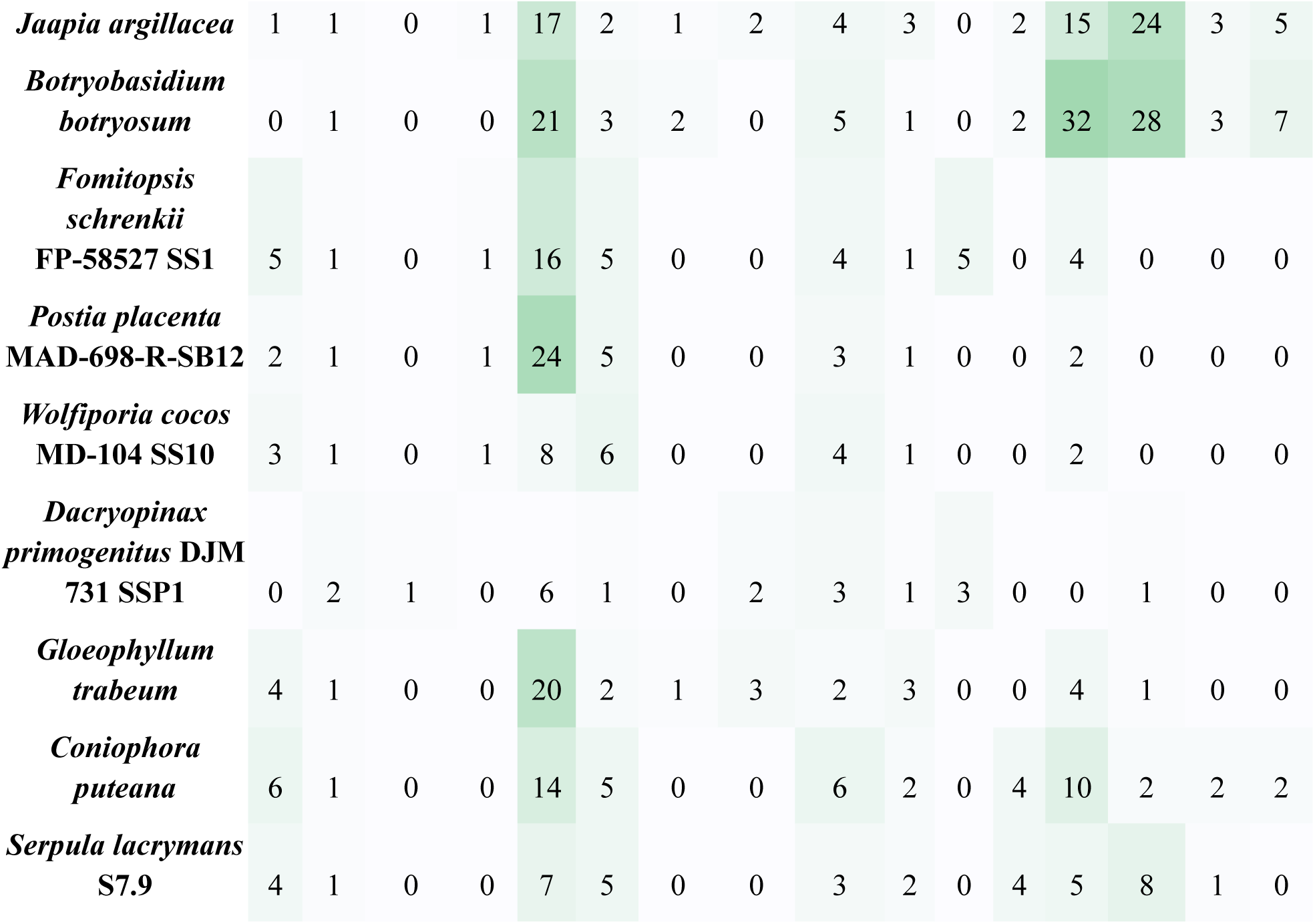
Counts of selected CAZyme families targeting crystalline cellulose and lignin, predicted from genomes of fungal species designated as white-rot (blue), brown-rot (brown), or uncertain (grey). Both *C. cesatii* (our genome of interest highlighted in yellow) and *C. variabilis* have multiple copies of AA9, CBM1, GH6, and GH7, but only single AA2 proteins suggesting deviation from white rot fungi towards uncertain types of wood decay.

Since according to CAZy the *C. cesatii* genome encoded only one AA2 encoding gene, we looked closer and used CUPP to predict three AA2-like proteins (Figure 2). The CUPP analysis confirmed the CAZy-predicted protein (ID 272474) as AA2:1.5 and identified two additional proteins with hypothetical peroxidase functions: AA2:10.1 (protein ID 1032398) and AA2:24.1 (protein ID 1043698). Importantly, *C. cesatii* does not encode AA2:1.1 (neon blue) commonly found in white-rot fungi. Instead, it encodes AA2:10.1 (dark blue) and AA2:24.1 (orange) which is broadly distributed across all fungi in our dataset, and AA2.1.5 (neon green), similar to three brown-rot and three white rot fungi. These findings show that *C. cesatii* AA2 profile is distinct from white-rot fungi although it shares some features with some white-rot and brown-rot fungi. Except for AA2:1.5, *C. variabilis* shows the same pattern as *C. cesatii*.

**Figure 2:**
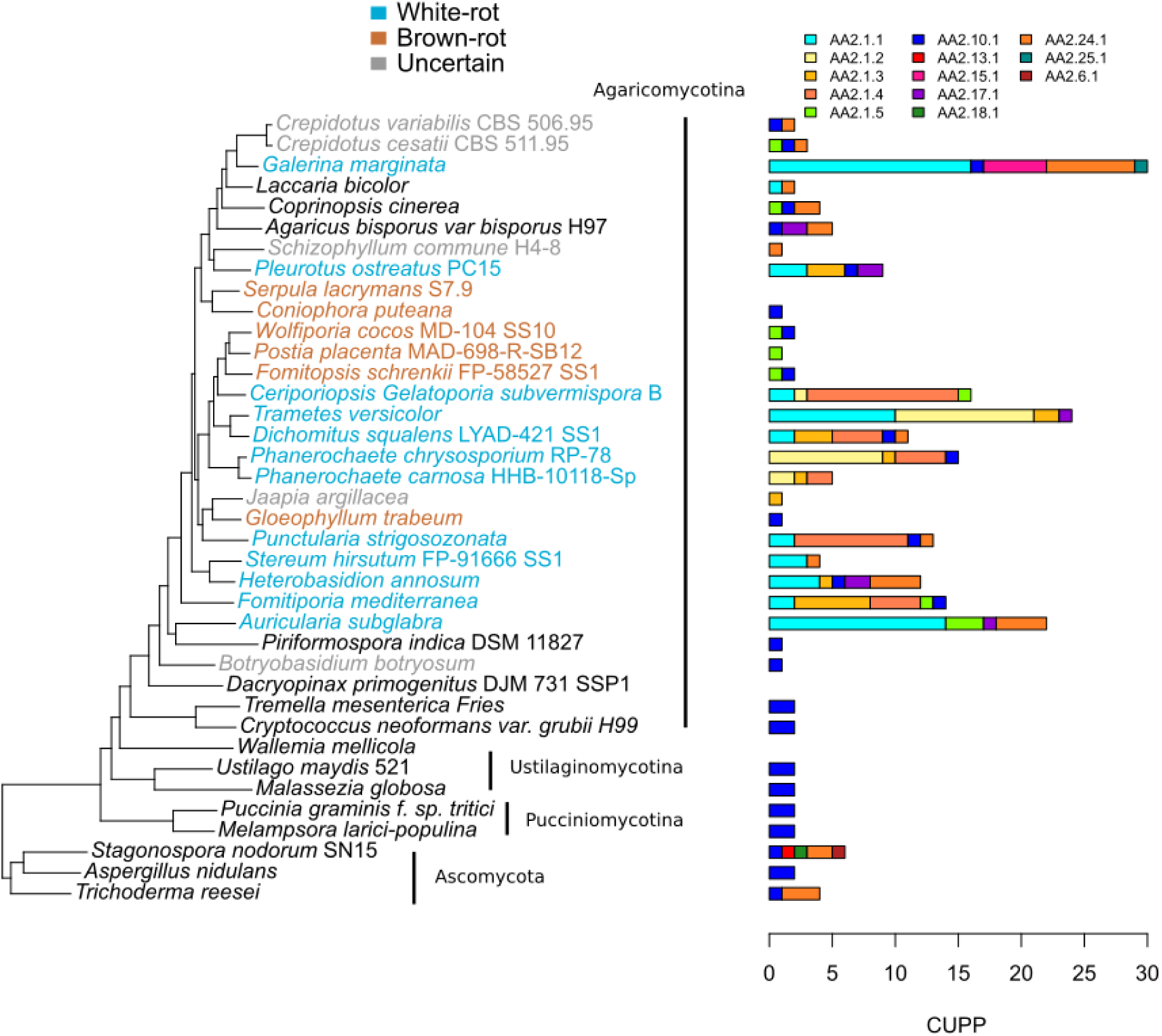
CUPP profile breakdown of predicted AA2 genes shows that *C. cesatii* has a pattern distinct from white-rot fungi and similar to brown-rot fungi (e.g. does not contain AA2.1.1 but contains AA2.1.5).

### Unique Orthogroup and PFAM Analysis

To further investigate the phylogenetic relationships between the genes, annotations, and protein classifications in white-rot fungi and brown-rot fungi, we examined OrthoFinder clusters for unique orthogroup patterns with notable functional properties. PFAM analysis provides additional insights into protein annotations, enabling us to compare orthogroups across species, their functions and evolutionary relationships.

Specifically, we sought examples where *C. cesatii* shared proteins with white-rot fungi, brown-rot fungi, and species with uncertain type of wood decay. Such analyses suggested orthogroups OG0004676 and OG0007022 as specific for white-rot fungi, and OG0000987 and OG0006907 specific for brown-rot fungi (Table 3). For the top two orthogroups for each group – white-rot, brown-rot, and uncertain-decay fungi - ring plots were generated (Figure 3) to show distribution of proteins across species and orthogroups. The white-rot cluster we focused on (from the ring plot) was filtered specifically to make white-rot comparisons. In other words, most of the white-rot fungi have at least one protein in this cluster whereas most of the brown-rot fungi lack proteins in this cluster. Figure 3 further shows that all of the white-rot fungi (including both the *Crepidotus* species) have PF02627 (Carboxymuconolactone decarboxylase family - see specific PFAM definitions in Table 4). Additionally, both *Crepidotus* species also have PF12697 (Alpha/beta hydrolase family) and PF12937 (F-box-like) which are not observed anywhere else in our dataset. Also, PF00734 is present in most white-rot fungi, fungi of uncertain type of decay, in *C. cesatii* but not in *C. variabilis*.

**Figure 3:**
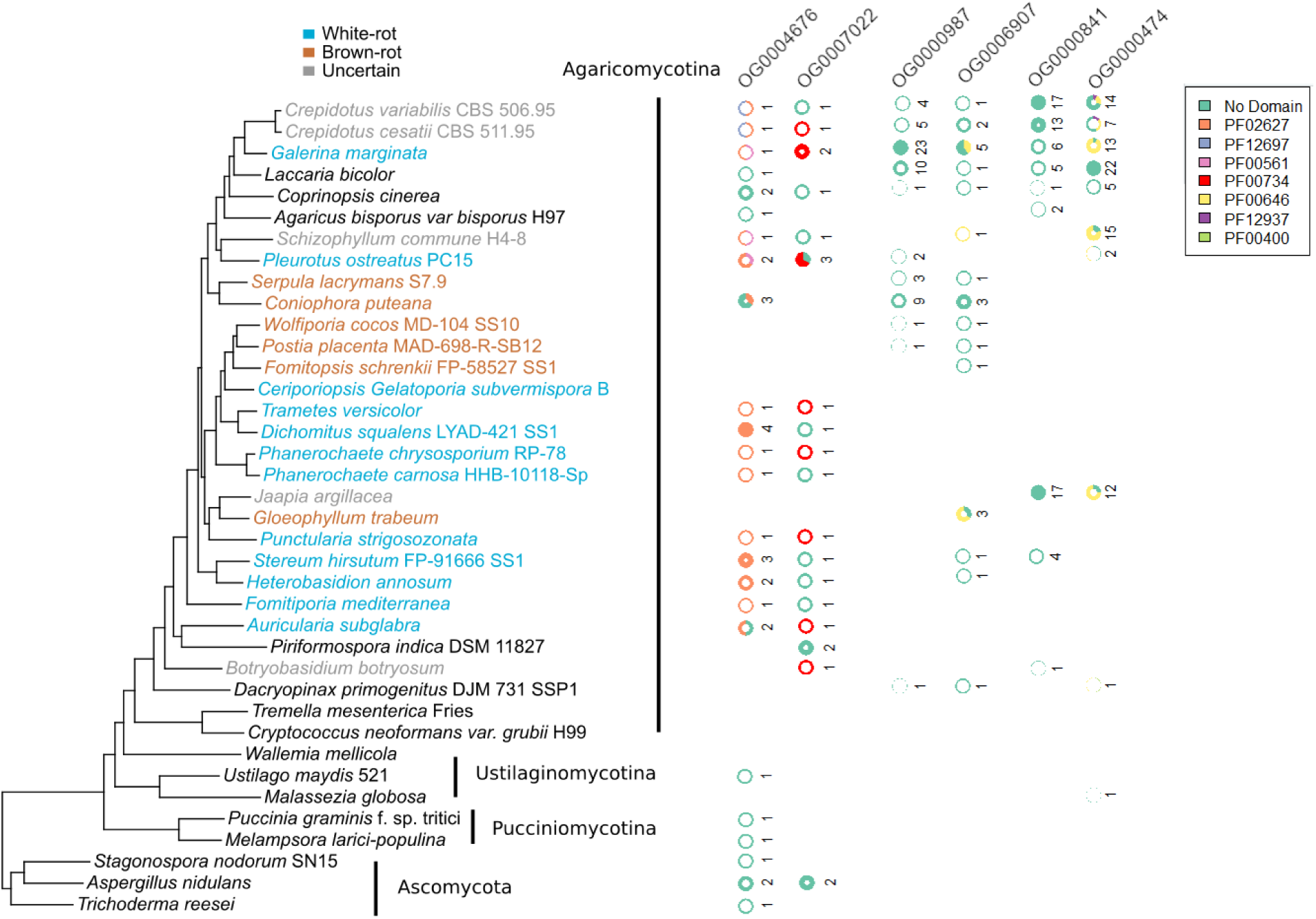
PFAM distributions within the top two orthogroups for each of the wood rot types. In the ring plot, the thickness of the "donut" indicates the relative size of the cluster, and the numbers represent the count of proteins from the organism in each cluster. PFAM Definitions with Orthogroup ID (if applicable) as follows: PF02627 – Carboxymuconolactone decarboxylase family – OG0004676; PF12697 – Alpha/beta hydrolase family – OG0004676; PF00561 – Alpha/beta hydrolase fold; PF00734 – Fungal cellulose binding domain – OG0007022; PF00646 – F-box domain; PF12937 – F-box-like – OG0000474; PF00400 – WD domain, G-beta repeat.

**Table 3:** Unique orthogroup specific to white-rot, brown-rot, and uncertain fungi, with PFAM annotations when available. The search allowed for flexibility in terms of presence within ingroups and absence from outgroups. By default, an orthogroup was identified if it was shared with at least 80% of the ingroup and missing from at least 80% of the outgroup. ^A^ Found in at least 70% of brown-rot and missing from at least 80% of non-brown-rot. ^B^ Found in at least 80% of brown-rot and missing from at least 70% of non-brown-rot fungi

| Ingroup Wood Decay Type | Orthogroup | *C. cesatii* PFAM Annotations |
| --- | --- | --- |
| White-Rot | OG0004676 | PF02627 Carboxymuconolactone decarboxylase family |
|  |  | PF12697 Alpha/beta hydrolase family |
| White-Rot | OG0007022 | PF00734 Fungal cellulose binding domain |
| Brown-Rot<sup>A</sup> | OG0000987 | No PFAM annotation for *C. cesatii* proteins |
| Brown-Rot<sup>B</sup> | OG0006907 | No PFAM annotation for *C. cesatii* proteins |
| Uncertain | OG000841 | No PFAM annotation for *C. cesatii* proteins |
| Uncertain | OG0000474 | PF00646 F-box domain |
|  |  | PF12937 F-box-like |
|  |  | PF00646 F-box domain |
| Uncertain | OG0006379 | No PFAM annotation for *C. cesatii* proteins |
| Uncertain | OG0001248 | PF07690 Major Facilitator Superfamily (X2) |
| Uncertain | OG0005130 | PF00850 Histone deacetylase domain |
| Uncertain | OG0001743 | No PFAM annotation for *C. cesatii* proteins |
| Uncertain | OG0008082 | PF05368 NmrA-like family (X2) |
| Uncertain | OG0008248 | PF13302 Acetyltransferase (GNAT) domain |
| Uncertain | OG0008437 | PF13086 AAA domain |
|  |  | PF13087 AAA domain |
| Uncertain | OG0009734 | PF00226 DnaJ domain |
|  |  | PF12796 Ankyrin repeats (3 copies) |
| Uncertain | OG0010638 | PF00172 Fungal Zn(2)-Cys(6) binuclear cluster domain |
| Uncertain | OG0008877 | No PFAM annotation for *C. cesatii* proteins |
| Uncertain | OG0006331 | PF00122 E1-E2 ATPase |
|  |  | PF00689 Cation transporting ATPase, C-terminus |
|  |  | PF00690 Cation transporter/ATPase, N-terminus |
|  |  | PF00702 haloacid dehalogenase-like hydrolase |
| Uncertain | OG0007188 | PF02417 Chromate transporter |
| Uncertain | OG0007870 | No PFAM annotation for *C. cesatii* proteins |
| Uncertain | OG0006877 | No PFAM annotation for *C. cesatii* proteins |
| Uncertain | OG0009743 | No PFAM annotation for *C. cesatii* proteins |
| Uncertain | OG0009730 | PF00067 Cytochrome P450 |
| Uncertain | OG0008329 | PF00646 F-box domain |
| Uncertain | OG0010291 | No PFAM annotation for *C. cesatii* proteins |
| Uncertain | OG0009727 | No PFAM annotation for *C. cesatii* proteins |
| Uncertain | OG0005793 | PF03211 Pectate lyase |
| Uncertain | OG0003757 | No PFAM annotation for *C. cesatii* proteins |
| Uncertain | OG0007610 | No PFAM annotation for *C. cesatii* proteins |
| Uncertain | OG0008678 | No PFAM annotation for *C. cesatii* proteins |
| Uncertain | OG0009300 | PF01753 MYND finger |
| Uncertain | OG0008891 | No PFAM annotation for *C. cesatii* proteins |

Taken together, these observations support the conclusion that *C. cesatii* has a PFAM distribution that is more similar to the white-rot fungi compared to the brown-rot fungi.

To summarize, *C. cesatii* is an example of wood-decaying fungi that shared some but not all characteristics of white-rot and brown-rot fungi, which illustrates a continuum of wood-decay fungi rather than a strict dichotomy. Specifically, PFAM and orthogroup patterns are more consistent with white-rot fungi while CUPP and CAZyme data show patterns consistent with white-rot fungi, except for AA2 and AA2.1.1, like in decay fungi of uncertain type. Thus, *C. cesatii* more likely represents an uncertain type of wood decay, not fully defined but characterized by a mosaic of white-rot and brown-rot features.

Comparison with its sister species *C. variabilis* adds further nuance. Prior studies have shown that *C. variabilis* can efficiently degrade the lipophilic compounds responsible for pitch deposition from eucalypt wood (Gutiérrez et al., 1999). However, *C. variabilis* has not been experimentally characterized in terms of type of wood decay. While initially *C. variabilis* was considered as part of the white-rot group, in the course of *C. cesatii* genome analysis both demonstrated most enzymes present in white rot fungi but lacking AA2 and more white-rot specific AA2.1.1 class II peroxidase. This suggests that both Crepidotus species may represent ‘intermediate’ type of wood decay, close to white rot but with a reduced complement of class II peroxidases.

Through the PFAM, orthogroups, CUPP, and CAZyme analysis, we identified *C. cesatii* as a potential wood decayer encoding enzymes such as carboxymuconolactone as well as domains of unknown function shared with other wood decayers of an uncertain rot type. Since the number of available fungal genomes from fungi with “uncertain” decay type is very limited, their deeper sequencing and molecular characterizations are critical to refine and expand classifications beyond the traditional white-rot/brown-rot framework.

## Conclusion

In conclusion, this comparative study places *C. cesatii* among fungi of the uncertain wood-decay type, supporting the emerging view that fungal decay strategies exist along a continuum rather than within a strict white rot and brown rot dichotomy. *C. cesatii* encodes multiple CAZymes typical of white rot fungi: CBM1 and AA9 which enhances hydrolysis of lignocellulose. However, based on CUPP analysis of AA2 (peroxidases), *C. cesatii* appears to have diverged from typical white-rot fungi by lacking to degrade lignin with class II peroxidases present in other white rot fungi. Furthermore, the PFAM distributions of identified orthogroups also suggested that *C. cesatii* has patterns similar to white-rot fungi compared to brown-rot fungi. Altogether, *C. cesatii* genome is similar to genomes of other fungi of uncertain type of wood decay, sharing some but not all characteristics of white-rot and brown-rot fungi, suggesting that *C. cesatii* may also represent an unknown, intermediate type of wood decay in a continuous spectrum between white-rot and brown-rot wood decayers. Taken together, these features suggest that both *Crepidotus* species*, C. cesatii* and *C. variabilis,* may not fit into the traditional white-rot/brown-rot dichotomy, mainly showing similarities to white rot fungi. This evidence further supports the notion that fungal decay strategies exist along a spectrum rather than as a strict binary classification (Riley et al., 2014).

Given these findings, future molecular studies are needed to clarify the classification of these fungi. Looking at the CAZymes and the CUPP analysis allows us to understand that some wood decaying fungi have different strategies compared to their white-rot and brown-rot counterparts. However, the current analysis methodologies are insufficient and more fungi need to be studied before a clear continuum can be established. Future studies should continue to focus on CAZyme and CUPP analysis while also taking into account functional nuances, post-translational modifications that can affect enzyme activity, and environmental factors. Both genome sequencing and molecular wood decay characterization for poorly characterized wood decay fungi will help better define the spectrum of fungi lifestyles. Through this, key insights into ecosystem function, agricultural sustainability, and biotechnological innovation to address global challenges can be achieved.

## Data Availability

Genome assembly and annotations are available from JGI fungal genome portal MycoCosm (https://mycocosm.jgi.doe.gov/Creces1) and have been deposited in GenBank under the accession no. JBQVAC000000000. The version described in this paper is version JBQVAC010000000.

## Contributions

● ST: Writing, Investigation
● SA: Annotation, Writing, Investigation, Supervision
● GH, KL, AL: Sequencing, Assembly
● DC and JM: DNA Extraction, Sample Preparation
● JS and KB: Project coordination
● IVG: Project coordination, Supervision, Editing

## Acknowledgments

The work (proposal 10.46936/10.25585/60001019) performed at the U.S. Department of Energy (DOE) US DOE Joint Genome Institute (https://ror.org/04xm1d337), a DOE Office of Science User Facility, is supported by the Office of Science of the U.S. Department of Energy operated under Contract No. DE-AC02-05CH11231.

